# Activity of lysosomal exoglycosidases in blood serum and urine of patients with tick-borne and other lymphocytic meningitis

**DOI:** 10.1101/2025.04.22.649922

**Authors:** Sławomir Dariusz Szajda, Jadwiga Snarska, Alina Kępka, Michalina KrzyŻak, Sławomir Pancewicz, Krzysztof Zwierz, Alina Minarowska

**Affiliations:** Department of Emergency Medical Service, University of Warmia and Mazury in Olsztyn, Olsztyn, Poland; Department of Surgery, Collegium Medicum University of Warmia and Mazury in Olsztyn, Olsztyn, Poland; Department of Biochemistry, Radioimmunology and Experimental Medicine, The Children’s Memorial Health Institute of Warsaw, Warsaw, Poland; Department of Hygiene, Epidemiology and Ergonomics, Medical University of Bialystok, Bialystok, Poland; Department of Infectious Diseases and Neuroinfection, Medical University of Bialystok, of Bialystok, Bialystok, Poland; Medical University of Bialystok, Bialystok, Poland; Chair of Pulmonology, School of Public Health, Collegium Medicum, University of Warmia and Mazury in Olsztyn, Poland; Outpatient Cystic Fibrosis Clinic, Department of Pediatrics, Gastroenterology, Hepatology, Nutrition, Allergology and Pulmonology, Children’s University Hospital in Bialystok, Poland

**Keywords:** tick-borne encephalitis (TBE), other (non-tick) lymphocytic meningitis (LMD), lysosomal exoglycosidases, blood serum, urine

## Abstract

Tick-borne encephalitis (TBE), and other (non-tick) lymphocytic meningitis diseases (LMD) are characterized by the presence of inflammatory processes differently located and with different intensity. Inflammatory processes in TBE and LMD may influence the activity of some catabolic enzymes including lysosomal exoglycosidases: N-acetyl-β-D-hexosaminidase (HEX), its isoenzymes A (HEX A) and B (HEX B) as well as β-galactosidase (GAL), α-mannosidase (MAN), α-fucosidase (FUC) and β-glucuronidase (GLU). The aim of the study was to investigate the activity of lysosomal exoglycosidases in the serum and urine of patients with TBE and LMD. Material and methods: Examined group included 41 patients: a/ 25 patients with TBE, b/ 16 patients with LMD. Control group included 25 patients in which Borrelia burgdoferi and inflammatory state in central nervous system were excluded. Lysosomal exoglycosidases activity were assayed in the serum and urine by the colorimetric method, with the p-nitrophenyl derivative of appropriate sugar as substrate. Results: Concentration of activity (pKat/mL) of HEX, HEX A, HEX B and GLU increases significantly in serum of patients with TBE, and LMD as well as GAL, MAN and FUC serum activity in LMD patients in comparison to serum of patients from the control group (p<0.01). A significant increase (p<0.05) in the specific activity of FUC (pKat/μg protein) and the activity of HEX, HEX B and GLU expressed in pKat/μg creatinine was also found in the urine of TBE patients compared to the control group. Our results suggests existence of negative correlation for GAL activity in serum and urine of patients with LMD (r=-0,568; p=0,022). Conclusions: 1. TBE and LMD change catabolism of glycoconjugates. 2. Lysosomal exoglycosidases activity concentrations in blood serum can be used in the diagnosis of TBE and LMD. 3. Concentrations activities of β-galactosidase, α-mannosidase and α-fucosidase in serum can vary from TBE, inflammation of other LMD. 4. Lysosomal exoglycosidase activity in urine converted to μg creatinine can distinguish TBE from LMD.

## INTRODUCTION

Lymphocytic meningitis and encephalitis are acute inflammatory diseases of the central nervous system (CNS). The most common etiologic agent of lymphocytic meningitis and encephalitis are viruses, mainly enteroviruses. Viruses can also cause lymphocytic meningitis and encephalitis: Herpes, cytomegalovirus, Epstein-Barr, parotitis, chickenpox, rubella, measles, tick-borne encephalitis, lymphocytic meningitis and vasculitis, influenza and paragroup, mycobacteria, Borrelia, Leptospira, Treponema, as well as protozoa and fungi [1]. Lymphocytic meningitis and encephalitis are usually diseases with a benign course and good prognosis. Above statement applies mainly to the well-studied pediatric population, with enteroviral infections and predominating chaperoned meningitis. In adults, meningitis and encephalitis of bacterial etiology (tuberculosis, neuroborreliosis) and severe viral infections of the central nervous system are observed more often than in children. Numerous diagnostic problems and limitations make it not always possible to determine the etiological agent of neuroinfections [1].

Tick-borne encephalitis (TBE) is a seasonal infectious disease caused by a Flavivirus of the Flaviviridae family. The main vectors and reservoirs of the TBE viruses are endemic ticks of the Ixodes family. Humans can become infected with the TBE viruses through the bite of an infected tick, and by drinking unpasteurized cow’s, sheep’s or goat’s milk. The course of TBE is biphasic: taking on the appearance of a flu-like infection in phase I and meningitis and encephalitis in phase II [1-3]. Factors influencing the course of TBE and the occurrence of complications include age, place of residence (TBE endemic areas), type of work, stress, past or concurrent illnesses associated with a weakened immune system and the coexistence of other infections, caused by other tick-borne pathogens (Borrelia burgdorferi, Anaplasma phagocytophilum). It is assumed that the percentage of neurological complications depends on the severity of the clinical course of the disease and affects 3-55% of patients. Frequent complications after a history of TBE are vegetative disorders, i.e. physical and mental fatigue, excessive nervous excitability, mood swings, explosiveness or excessive sweating. Serious complications after a history of TBE are disorders of the intellectual and mental spheres [1].

The main etiologic agents of other (non-tick) lymphocytic meningitis (LMD) are enteroviruses. Herpes, cytomegalovirus, Epstein-Barr, parotitis, chicken pox, rubella, measles, lymphocytic meningitis and choroid plexus, influenza as well as paragroup viruses can also cause LMD. Mycobacteria, Leptospira, Treponema, protozoa and fungi can also cause lymphocytic meningitis [1].

Inflammatory processes in tick-borne encephalitis and lymphocytic meningitis can affect the activity of certain catabolic enzymes, including lysosomal exoglycosidases: N-acetyl-β-D-hexosaminidase (HEX), its isoenzymes A (HEX A) and B (HEX B), as well as β-galactosidase (GAL), α-mannosidase (MAN), α-fucosidase (FUC) and β-glucuronidase (GLU). These are lysosomal hydrolases that are involved in the hydrolysis of oligosaccharide chains of glycoconjugates: glycoproteins, glycosaminoglycans and glycolipids [4].

Thea im of the paper was determination the activity of lysosomal exoglycosidases in serum and urine of the patients with TBE and LMD of different etiology and evaluation of: 1. Glycoconjugate catabolism in different types of lymphocytic meningitis. 2. Differences in activity of lysosomal exoglycosidases: HEX and its isoenzymes HEX A and HEX B, MAN, FUC, GLU in serum and urine of patients with different types of lymphocytic meningitis. 3. Usefullness the determination of lysosomal exoglycosidases in diagnostics of different types of lymphocytic meningitis.

## MATERIAL AND METODS

The study was conducted in blood serum and urine remaining after routine diagnostic laboratory tests. The study group consisted of 41 patients treated at the Department of Infectious Diseases and Neuroinfection, Medical University of Bialystok: a/ 25 patients with tick-borne encephalitis (12 women, 13 men) aged 21 to 71 years, mean age 46.04 ±14.70 years, b/ 16 patients with lymphocytic meningitis of non-tick etiology (8 women, 8 men) aged 24 to 67 years, mean age 41.94 ±15.15 years. The diagnosis was based on epidemiological history, clinical picture of the disease, results of laboratory tests mainly CSF examination and serological tests. The control group consisted of 25 subjects (8 women, 17 men) aged 18 to 74 years, mean age 33.20 ±14.15, in whom Borrelia burgdorferi infection and the presence of inflammation within the CNS were excluded. Diabetes mellitus, obesity, alcoholism, kidney and liver diseases were also exclusion criteria for the study group.

Examination the activity of lysosomal exoglycosidases: HEX, its isoenzymes HEX A and HEX B, GAL, MAN, FUC and GLU was conducted in double trials using colorimetric method of Chatterjee et al [5], modified by Zwierz et al [6] and adapted to assays on microplates by Marciniak et al [7]. Concentration of lysosomal exoglycosidase activity was expressed in pKat/mL, specific activity in pKat/µg protein, and activity per µg creatinine in pKat/µg creatinine. Total protein concentration was determined using the “ diagnostic reagent for in vitro quantification of total protein concentration in urine” with pyrogallol red, using a ready-made reagent from ABX Pentra and a Pentra 400 biochemical analyzer from Horiba ABX SAS, Montphellier, France [8]. Creatinine concentration was determined using the “ diagnostic reagent for in vitro quantification of creatinine concentration in serum, plasma and urine by a colorimetric method” based on the reaction of creatinine with alkaline picrate, using a ready-made reagent from ABX Penrta and a Pentra 400 biochemical analyzer from Horiba ABX SAS, Montphellier, France [9].

In statistical analysis, normality was verified using Kolmogorov-Smirnov test improved by Lillefors and Shapiro-Wilk test. Nonparametric ANOVA test for Kruskal-Wallis ranks and post-hoc test with the possibility of multiple comparison of ranks for every trial in case of many groups were applied in order to compare quantitative variables, excluding normality. The Wilcoxon test (matched-pairs) was applied in order to compare dependable variables in case of two variables. Spearman’s rank correlation coefficient was also assigned. Level of statistical significance was established at p<0.05.

## RESULTS

The study shows that serum concentrations of HEX activity in TBE patients [Me = 1360.74 (Q1 = 1068.33; Q2 = 1811.29)] and LMD [Me = 1429.32 (Q1 = 866.26; Q2 = 1686.29)] were more than three times, significantly higher (p<0.001) than serum HEX activity in the control subjects [Me = 413.38 (Q1 - 317.02; Q2 = 448.61)]. Comparison of serum HEX activity concentrations in TBE and LMD patients shows that there were no significant differences between the groups (Figure 1).

**Figure 1.**
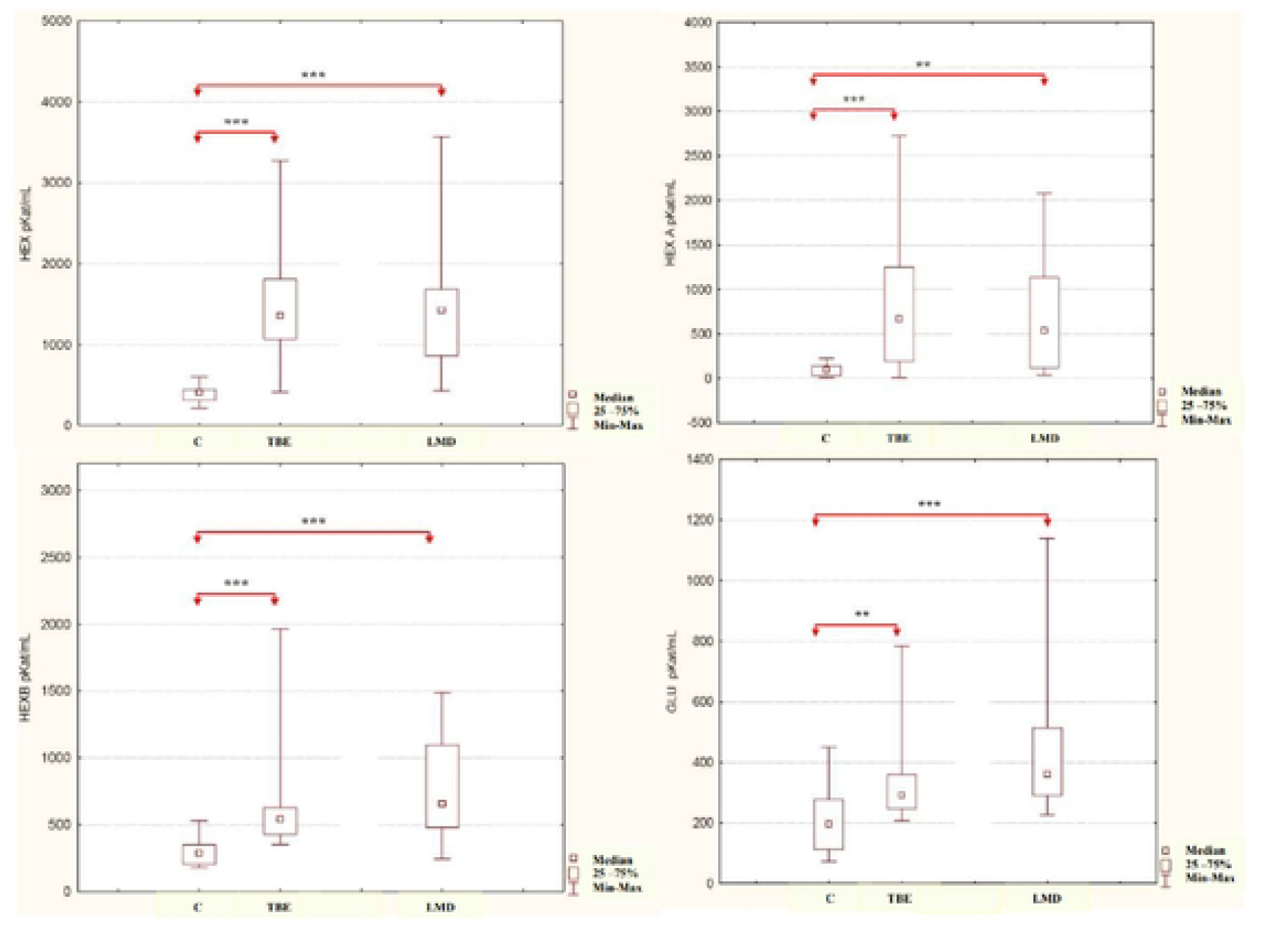
Activity concentrations (pKat/mL) of HEX, HEX A, HEX B and GLU in serum of patients with tick-borne encephalitis (TBE), other lymphocytic meningitis (LMD) and controls (C): ** p < 0.01, *** p < 0.001.

The serum concentration of HEX A activity in TBE patients [Me = 668.61 (Q1 = 194.67; Q2 = 1252.61)] was more than six and a half times higher (p<0.001), and in LMD patients [Me = 535. 35 (Q1 = 120.00; Q2 = 1134.20)] was more than five times higher (p<0.01) compared to the serum HEX A activity concentration of control subjects [Me = 100.37 (Q1 = 28.58; Q2 = 140.02)]. The serum concentration of HEX A activity in TBE patients tended to increase as compared to the serum concentration of HEX A activity in LMD patients (Figure 1).

The serum concentration of HEX B activity in TBE patients [Me = 539.02 (Q1 = 424.74; Q2 = 626.37)] was almost twice as high (p<0.001), and in LMD patients [Me = 656.20 (Q1 = 477. 96; Q2 = 1094.59)] was more than twice as high (p<0.001) as compared to serum HEX B activity levels in the control subjects [Me = 286.85 (Q1 = 204.35; Q2 = 346.87)]. Serum concentrations of HEX B activity in TBE and LMD patients showed no significant differences between the compared groups (Figure 1).

In contrast, serum GLU activity levels in TBE patients [Me = 291.86 (Q1 = 247.57; Q2 = 360.44)] were significantly almost one and a half times higher (p<0.01), and in LMD patients [Me = 360. 97 (Q1 = 290.97; Q2 = 514.39)] was significantly almost two times higher (p<0.001) as compared to the serum GLU activity concentration of control subjects [ME = 196.04 (Q1 = 113.33; Q2 = 277.57)]. The serum GLU activity concentration of TBE patients tended to decrease as compared to the serum GLU activity concentration of LMD patients (Figure 1).

The results show that the serum concentration of GAL activity in LMD patients [Me = 246.14 (Q1 = 125.59; Q2 = 478.11)] was more than two and a half times higher (p<0.001) than the serum concentration of GLU activity in control subjects [Me = 94.66 (Q1 = 72.16; Q2 = 130.77)]. The serum concentration of GLA activity in LMD patients was almost two and a half times higher than the serum concentration of activity in TBE patients [Me = 98.98 (Q1 = 86.47; Q2 = 155.41)] and indicated a strong trend for significant differences between TBE and LMD (statistical significance slightly above the cutoff value of p = 0.072). The serum concentration of GAL activity in TBE patients tended to increase as compared to the serum concentration of GLA activity in the control subjects (Figure 2).

**Figure 2.**
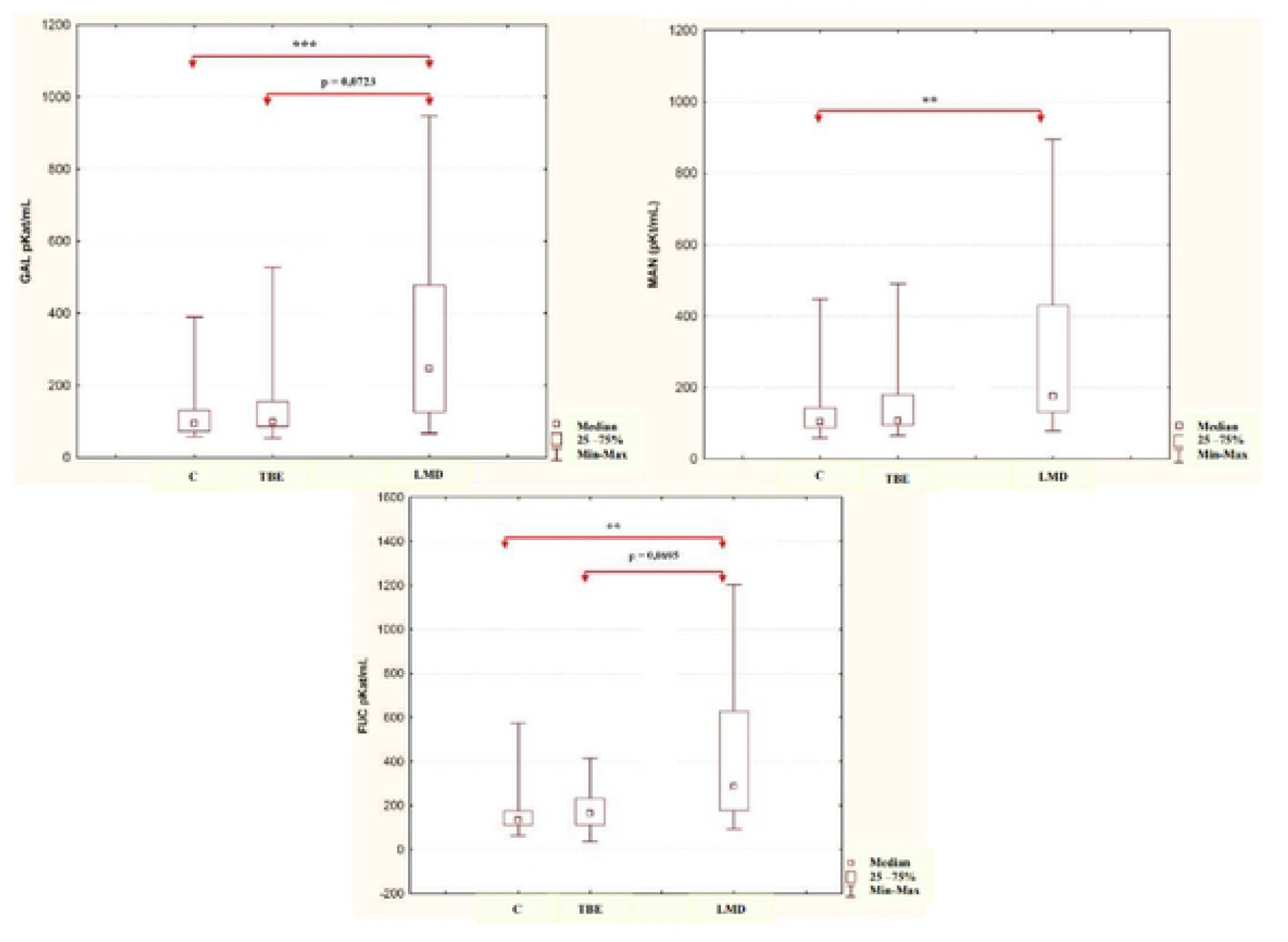
Activity concentrations (pKat/mL) of GAL, MAN and FUC in serum of patients with tick-borne encephalitis (TBE), other lymphocytic meningitis (LMD) and controls (C): ** p < 0.01, *** p < 0.001.

The serum concentration of MAN activity in LMD patients [Me = 175.59 (Q1 = 131.03; Q2 = 429.88)] was more than one and a half times higher (p<0.01) as compared to the serum concentration of MAN activity in control subjects [Me = 103.80 (Q1 = 87.35; Q2 = 142.91)]. The mean serum MAN activity concentration of LMD patients was more than one and a half times higher than the serum activity concentration of TBE patients [Me = 107.55 (Q1 = 95.05; Q2 = 179.70)], but indicated only an upward trend. The serum concentration of MAN activity in TBE patients tended to increase compared to the serum concentration of MAN activity in control subjects (Figure 2).

In contrast, the serum FUC activity concentration of LMD patients [Me = 287.22 (Q1 = 175.59; Q2 = 628.86)] was more than twice as high (p<0.01) as the serum FUC activity concentration of control subjects [Me = 135.41 (Q1 = 110.75; Q2 = 175.06)]. The serum concentration of FUC activity in LMD patients was almost twice as high as the serum concentration of activity in TBE patients [Me = 166.13 (Q1 = 111.12; Q2 = 231.49)] and showed a strong upward trend (statistical significance slightly above the cutoff value of p = 0.0696). The serum FUC activity concentration of TBE patients had a weak upward trend compared to the serum FUC activity concentration of the control subjects (Figure 2).

Urine analysis showed no significant differences in the activity levels of HEX, its isoenzymes A and B, GAL, MAN, FUC, and GLU in TBE and LMD patients as compared to the urine levels of the control subjects (Table 1).

In contrast, it was shown that the specific activity of FUC in the urine of TBE patients [Me = 3.27 (Q1 = 1.62; Q2 = 4.76)] was more than one and a half times lower (p<0.05) as compared to the specific activity of FUC in the urine of control subjects [Me = 5.44 (Q1 = 3.94; Q2 = 7.25)]. The specific activity of FUC in the urine of LMD patients [Me = 4.64 (Q1 = 2.39; Q2 = 5.96)] tended to decrease as compared to the specific activity of FUC in the urine of control subjects. The specific activity of FUC in the urine of LMD patients tended to increase as compared to the specific activity of FUC in the urine of TBE patients (Figure 3). Table 2 shows that there were no significant differences in the specific activity of HEX, its isoenzymes A and B, GAL, MAN and GLU in the urine of TBE and LMD patients as compared to the specific activity in the urine of control subjects.

**Figure 3.**
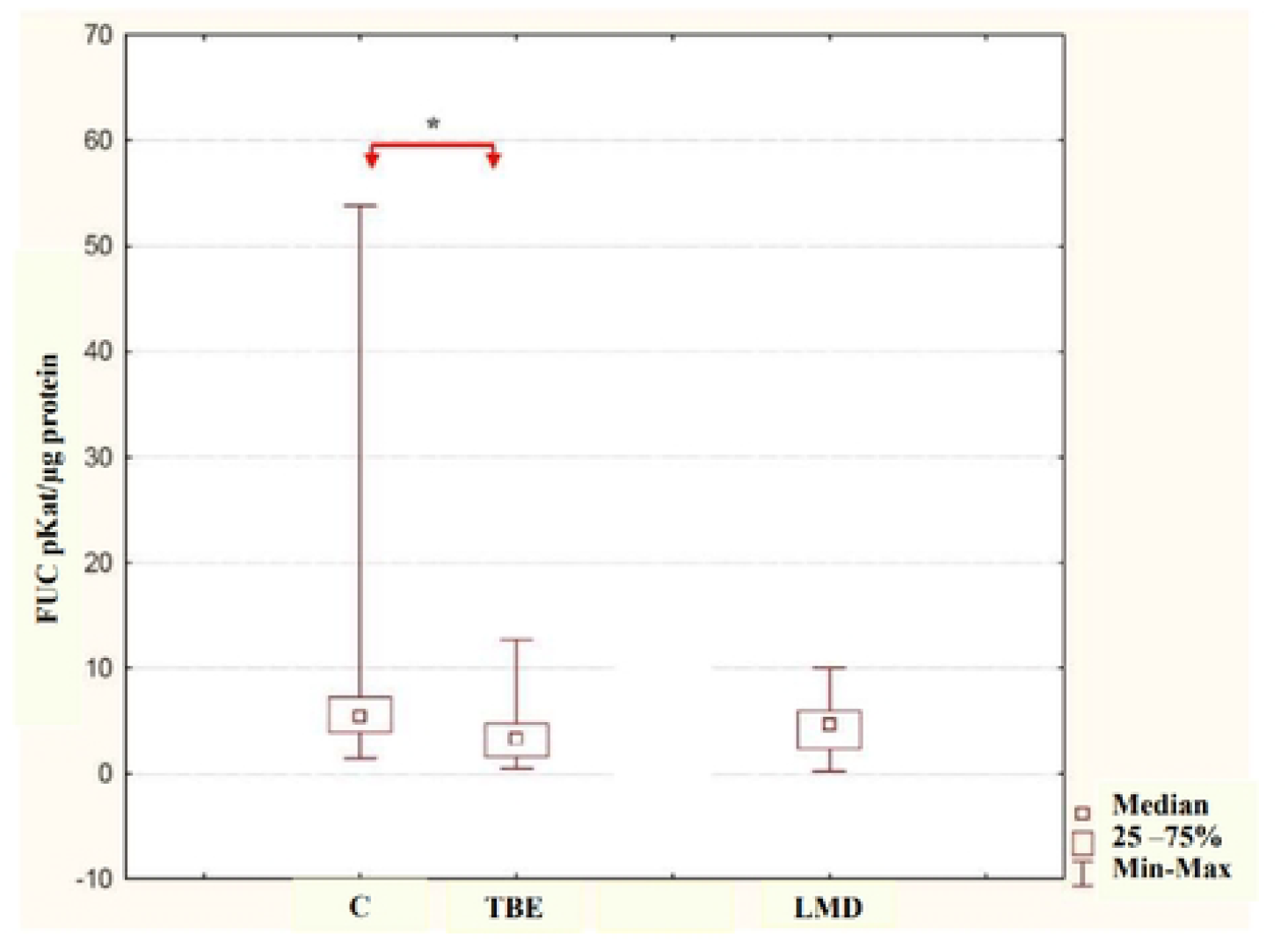
Specific activity (pKat/µg protein) of FUC in the urine of patients with tick-borne encephalitis (TBE), other lymphocytic meningitis (LMD), and controls (C): * p < 0.05.

In addition, we found that HEX activity per µg of creatinine in the urine of TBE patients [Me = 0.40 (Q1 = 0.31; Q2 = 1.27)] was significantly, more than three times, higher (p<0.05) than HEX activity in the urine of control subjects [Me = 0.13 (Q1 = 0.07; Q2 = 0.47)]. HEX activity per µg of creatinine in the urine of LMD patients [Me = 0.37 (Q1 = 0.12; Q2 = 1.28)] was almost three times higher than HEX activity in the urine of control subjects but showed no significant differences. HEX activity per µg of creatinine in TBE patients had a slight tendency to increase as compared to LMD patients (Figure 4).

**Figure 4.**
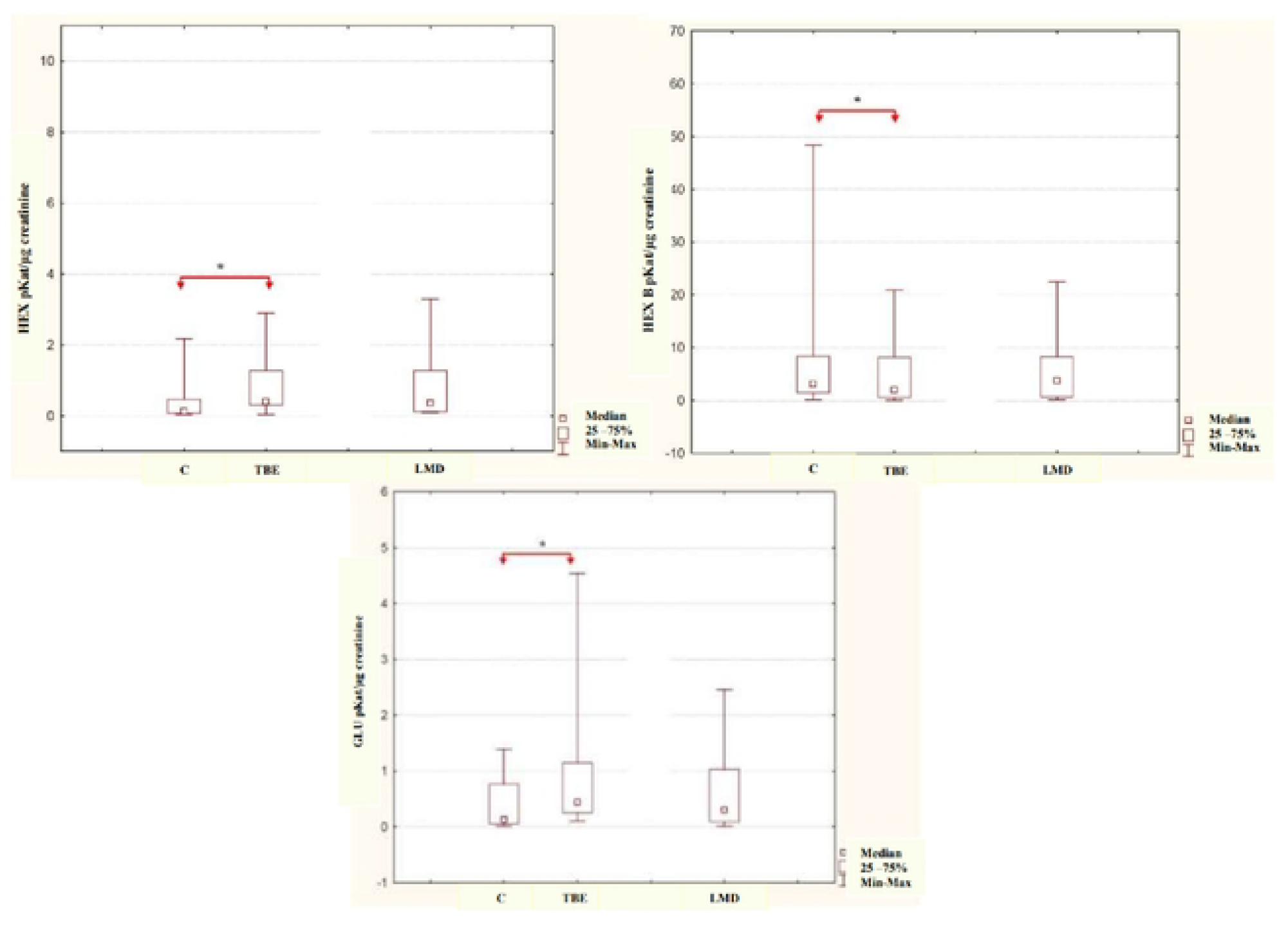
HEX, HEX B and GLU activities expressed per µg creatinine (pKat/µg creatinine) in the urine of patients with tick-borne encephalitis (TBE), other lymphocytic meningitis (LMD) and controls (C): * p < 0.05.

Also, HEX B activity expressed per µg creatinine in the urine of TBE patients [Me = 0.30 (Q1 = 0.16; Q2 = 0.70)] was significantly, more than four times higher (p<0.05) than HEX B activity in the urine of control subjects [Me = 0.07 (Q1 = 0.05; Q2 = 0.33)]. HEX B activity expressed per µg of creatinine in the urine of LMD patients [Me = 0.15 (Q1 = 0.10; Q2 = 0.42)] was non-significantly, more than twice as high as HEX B activity in the urine of control subjects. HEX B activity expressed per µg creatinine in the urine of TBE patients tended to decrease as compared to HEX B activity in the urine of LMD patients (Figure 4).

The GLU activity expressed per µg of creatinine in the urine of TBE patients [Me = 0.44 (Q1 = 0.26; Q2 = 1.15)] was found to be significantly, more than three times higher (p<0.05) than the GLU activity in the urine of control subjects [Me = 0.13 (Q1 = 0.06; Q2 = 0.77)]. GLU activity expressed per µg of creatinine in the urine of LMD patients [Me = 0.31 (Q1 = 0.09; Q2 = 1.03)] is non-significantly, more than twice as high as GLU activity in the urine of control subjects. GLU activity per µg of creatinine in the urine of TBE patients tends to increase as compared to that in the urine of LMD patients (Figure 4).

Analysis of the evaluated activities of the lysosomal exoglycosidases shows that GAL activity per µg of creatinine in the urine of TBE patients has a strong tendency to increase as compared to GAL activity in the urine of control subjects. HEX A, MAN and FUC activities per µg of creatinine in the urine of TBE and LMD patients and GAL in LMD tend to increase as compared to GAL activity in the urine of control subjects (Table 3).

Urine protein and creatinine testing showed a slight increase in protein and decrease in creatinine in TBE patients (Table 4).

The study suggests negative correlations (correlations) for: GAL activity concentration in serum and urine of LMD patients (r=-0.568 p=0.022); and a strong tendency to correlate GLU activity concentration in serum and urine of LM patients (r=-0.476; p=0.062).

## DISCUSSION

Lymphocytic meningitis and encephalitis are acute inflammatory diseases of the central nervous system (CNS) [1]. The most common etiologic agent of lymphocytic meningitis and encephalitis are viruses, mainly enteroviruses. Lymphocytic meningitis and encephalitis can also be caused by tick-borne encephalitis virus and Borrelia spirochetes [1]. One of the main sources of cases of lymphocytic meningitis and encephalitis are ticks. They transmit numerous pathogens that cause infectious diseases: viral, bacterial, parasitic and fungal. Diseases transmitted exclusively by ticks are tick-borne encephalitis and meningitis, Lyme disease (Lyme disease), granulocytic anaplasmosis (ehrlichiosis) and babesiosis [1]. The most common tick-borne diseases in Europe are Lyme borreliosis (BL) and tick-borne encephalitis (TBE) [1,10]. TBE is an acute viral infectious disease that occurs in various clinical forms, mainly affecting the central nervous system [11]. TBE is characterized by the presence of an inflammatory process of varying localization and intensity [2,3,11-13]. The cause of the inflammatory process in TBE is a virus of the Flaviviridae family and the genus Flavivirus. In TBE, the following cytokines are responsible for the development of the inflammatory process: tumor necrosis factor (TNF)-alpha, interleukins (IL) 1, 6 and 8 [14].The TBE virus multiplies in cells at the site of infection. It then spreads through the pathway of lymphatic vessels to the surrounding lymph nodes and liver, spleen, bone marrow in which it multiplies further. During the incubation period of the virus, which lasts from 2 to 28 days, no clinical symptoms are observed [1-3,11]. The virus incubation period is followed by the first phase of TBE, lasting 1 to 5 days. During the first phase of TBE, there is an increase in body temperature and the onset of nonspecific flu-like symptoms. After the period of initial symptoms, i.e. 3 weeks after the tick bite, the second phase of TBE - neurological - takes place. In the second phase of the disease, the TBE virus penetrates the CNS. The presence and replication of the virus in the CNS leads to: acute inflammation of the meninges and brain, congestion, appearance of bloody petechiae, inflammatory infiltrates, necrosis of microglia cells and cellular dysfunction. Inflammatory lesions are localized mainly in the periventricular area and cerebellum. The necrotic changes observed in TBE are similar to those in ischemic brain injury. In the second phase of TBE, there is a renewed sudden increase in body temperature reaching 40°C, severe headache, nausea, vomiting, meningeal symptoms, muscle and joint pain [3,11,12]. In both TBE and other tick-borne lymphocytic meningitis (LMD), damage to nervous system structures is associated with direct invasion by the pathogen on the one hand, and on the other hand is the result of inflammatory processes and toxic-metabolic mechanisms developing in response to infection [1,3]. Inflammatory processes taking place in the course of TBE and LMD can affect the activity of certain catabolic enzymes including lysosomal exoglycosidases. For example, β-glucuronidase in inflammation is released from granulocytes [15]. Serum levels of pro-inflammatory cytokines, including interleukin 1 (IL-1) and c-reactive protein (CRP), correlate very well with β-glucuronidase activity in the serum of patients with immune disorders [16,17]. Under moderate oxidative stress, rupture of some lysosomes is observed, accompanied by progressive apoptosis and further loss of intact lysosomes. The efflux of lysosomal enzymes into the cytoplasm is an important process accompanying apoptosis [16]. Lysosomal exoglycosidases: N-acetyl-β-D-hexosaminidase (HEX), its isoenzymes A (HEX A) and B (HEX B), β-galactosidase (GAL), α-mannosidase (MAN), α-fucosidase (FUC) and β-glucuronidase (GLU) are glycoproteins belonging to the lysosomal hydrolases. They take part in the hydrolysis of the sugar chains of glycoconjugates (glycoproteins, glycosaminoglycans and glycolipids) [4]. Glycoconjugates play an important role in the structure and function of both normal and pathological tissues [18,19]. They are involved in many cellular and molecular processes t.i.e.: cell differentiation, proliferation and growth, cell-to-cell interaction and cell-to-cell signal transduction [20]. Glycolipids and glycoproteins are responsible for the structure and function of neurons and glial cells in the brain. Some of the gangliosides and proteoglycans help regenerate damaged neurons and stimulate their growth [21,22]. Glycoconjugates (glycoproteins, proteoglycans and glycosaminoglycans: hyaluronic acid, chondroitin sulfates, dermatan, heparin) are essential components of the extracellular matrix (ECM) and basement membranes (BM) [23]. Changes in the structure of the ECM and cell membrane components can be crucial in many pathological processes [24,25]. Modification or destruction of ECM proteins by cancer cells, for example, can occur due to the release of enzymes that solubilize ECM components [26,27].The presence of lysosomal exoglycosidases has been demonstrated in many organs: liver, brain, kidney, stomach and biological fluids: blood, urine, cerebrospinal fluid, joint fluid and saliva [28]. In reviewing the literature, we found no publications dealing with the activity of lysosomal exoglycosidases catabolizing glycoconjugates in BTE or LMD. A study by Wielgat et al [20] demonstrates the presence of lysosomal exoglycosidases: HEX, HEX A and HEX B, GAL and MAN in brain tissues. They showed a significant increase in the activity of lysosomal exoglycosidases in glioblastoma malignancies compared to healthy brain tissues and non-cancerous brain glial cell lesions. According to Wielgat et al, lysosomal exoglycosidases [20] may be involved in the development of glial brain tumors. Our own study shows that the activity levels (pKat/mL) of lysosomal exoglycosidases: HEX, HEX A, HEX B and GLU significantly increase in the serum of BTE and LMD patients compared to the serum of control subjects (Figure 1). We found a significant increase in GAL, MAN and FUC activities in the blood serum of LMD patients as compared to the blood serum of control subjects (Figure 2).The absence of a significant increase in serum GAL, MAN and FUC activities in BTE patients suggests the possibility of using the determination of these enzymes in differentiating LMD from BTE (Figure 2). We demonstrated an increase in the serum levels of GAL and FUC activity slightly above the significance limit in BTE patients as compared to LMD (Figure 2) with no significant differences in the activity of lysosomal exoglycosidases between the other compared groups of BTE patients, LMD and controls. It seems that determination of serum GAL and FUC activity levels could be applicable in differentiating BTE from LMD. However, such a finding needs to be confirmed in studies on a larger group of patients with BTE and LMD. The reason for the increase in serum levels of lysosomal exoglycosidases activity in patients with BTE or LMD could be the ongoing inflammatory process in the brain [1,3,12]. Disruption of the normal functioning of the blood-brain barrier observed in BTE [2] may also cause the passage of lysosomal exoglycosidases from the brain into the bloodstream and thus contribute to their increased activity in the serum of patients. Elevated activity of lysosomal exoglycosidases in blood serum indicates their active participation in the remodeling of extracellular matrix (ECM) glycoconjugates and basement membranes of the human body. Pancewicz et al [29] examined the activity of lysosomal exoglycosidases: HEX, GAL, MAN, FUC and GLU in cerebrospinal fluid collected from BTE patients before and two weeks after treatment. Their study shows that the activity of lysosomal exoglycosidases in the CSF before BTE treatment was significantly lower than in the control group. After BTE treatment and normalization of CSF parameters, the activity of HEX increased significantly, while the activities of MAN, GAL FUC and GLU showed an increasing trend after BTE treatment and were lower than the activities of the tested lysosomal exoglycosidases in the CSF of the control group. Perez et al [30] examined the activity of HEX and its isoenzymes in the CSF and blood serum of patients with multiple sclerosis. They found significantly higher activity of HEX and its isoenzymes in the cerebrospinal fluid of patients with multiple sclerosis compared to controls. In blood serum, they showed no significant differences in HEX activity between the control group and people with multiple sclerosis. In recent years, there have been major scientific advances explaining the relationship between structural changes in cellular organelles and biochemical metabolic disorders. Attention has been drawn to the use of non-invasive methods with high sensitivity to determine functional renal function [31]. Therefore, one of the goals of our work became the analysis of changes in the activity of lysosomal exoglycosidases in the urine of patients with BTE and LMD. Elevated levels of total protein in the urine (proteinuria) are detected in most renal diseases. Primary and secondary nephropathy can cause increased glomerular filtration or decreased protein reabsorption in the renal tubules. Extra-renal factors that cause proteinuria include infections, bleeding and urinary tract malignancies. Elevated urinary protein concentrations can also be associated with other acute illnesses, such as fever, also with exercise and mental stress [8]. Urinary creatinine concentration significantly decreases during starvation, as well as in acute and chronic renal failure. Creatinine is secreted from the body only by the kidneys, and is not reabsorbed or secreted by renal tubule cells. Creatinine serum concentration negatively correlates with glomerular filtration rate by which creatinine is considered a useful marker of renal filtration function [31,32]. Examination of protein and creatinine concentrations in urine allowed us to find, slightly above the limit of statistical significance, an increase in protein concentration and a decrease in creatinine concentration in BTE patients compared to the urine of healthy subjects (Table 4), which may be indicative of renal dysfunction that may be mediated by BTE. Renal glomeruli do not permeate proteins with a molecular weight greater than 68 kD [33], so serum lysosomal exoglycosidases are not filtered through a properly functioning renal tubule filtration membrane. However, trace amounts of HEX are detectable in physiological urine, indicating that it is not only cell breakdown and cell membrane damage that releases this enzyme. The appearance of HEX activity in urine may be due to the natural exfoliation of the renal tubule epithelium. Another reason for the presence of HEX in physiological urine may be the escape of lysosomal enzymes into the cytoplasm and outside the cell, and in the case of renal tubules into the urine [34,35] under conditions of normal cell metabolism. In our study, we found no significant differences between the activity levels (pKat/mL) of lysosomal exoglycosidases in the urine of healthy subjects and patients with BTE and LMD (Table 1), which may indicate the absence of renal damage by pathogens causing BTE and LMD. In order to exclude the influence of the amount of consumed fluids on the activity of the tested lysosomal exoglycosidases, we decided to convert the concentration of protein and creatinine in urine into specific activity (pKat/µg protein) and activity per µg of creatinine (pKat/µg creatinine). Due to the possibility of a relationship between enzyme activity per unit volume and the amount of minute diuresis, it is assumed that the best way to eliminate the effect of the amount of diuresis on urinary enzyme activity is to relate it to the concentration of creatinine in the urine [31,32]. By examining the specific activity (pKat/µg protein) of lysosomal exoglycosidases, we found that BTE significantly decreases the specific activity of FUC in the urine of patients compared to the urine of the control group (Figure 3) with no significant differences in the specific activity of other lysosomal exoglycosidases between the compared groups (Table 2), which may suggest that BTE affects renal function. In addition, we found a significant increase in the activity per µg of creatinine of HEX, HEX B and GLU in the urine of BTE patients compared to the activity of the above enzymes in the urine of control subjects (Figure 4). We also observed a trend toward an increase, slightly above the significance limit, of GAL in the urine of BTE patients compared to the activity per µg of creatinine in the urine of control subjects (Table 3). HEX, HEX B, GAL and GLU activities per µg creatinin were not significantly increased in LMD patients (Figure 4, Table 3). We observed no significant increase in HEX A, MAN and FUC activity per µg creatinin in BTE and LMD patients compared to the urine of control subjects (Table 3). It seems that the determination of activity per µg creatinine of HEX, HEX B and GLU or GAL in urine may be applicable in differentiating BTE from LMD. Examination of GAL and GLU activity levels in LMD patients indicates that there are negative correlations between serum and urine activity of these enzymes in the groups studied, suggesting that these enzymes do not pass from blood to urine. HEX is the most active of the lysosomal exoglycosidases studied. Studies by Szajda et al [36] suggest that the determination of HEX, HEX A and HEX B activity in blood serum may have applications in the early diagnosis of colorectal cancer. Assuming that exoglycosidases can act as an early indicator of the inflammatory process, it seems appropriate to consider their use in the early diagnosis of LMD, particularly TBE.

## CONCLUSIONS

The study conducted jumps to the following conclusions:

1. Tick-borne encephalitis and non-tick lymphocytic meningitis alter glycoconjugate catabolism.
2. Serum concentrations of lysosomal exoglycosidases activity can be used in the diagnosis of tick-borne encephalitis and other-than-tick lymphocytic meningitis.
3. Serum concentrations of β-galactosidase, α-mannosidase and α-fucosidase activities can differentiate tick-borne encephalitis from other lymphocytic meningitis.
4. Urinary lysosomal exoglycosidases activities converted to µg of creatinine can differentiate tick-borne encephalitis from other tick-borne meningitis.

## Author Contributions

## Funding

This research received no external funding.

## Institutional Review Board Statement

The study was conducted in accordance with the Declaration of Helsinki, and approved by the Ethics Committee of the

Medical University of Bialystok no: R-I-002/75/2012.

## Data Availability Statement

The original contributions presented in the study are included in the article; further inquiries can be directed to the corresponding author.

## Conflicts of Interest

The authors declare no competing interest.

## List of figures

## List of tables

**Table 1.** Activity concentrations (pKat/mL) of lysosomal exoglycosidases (HEX, HEX A, HEX B, GAL, MAN, FUC and GLU) in the urine of patients with tick-borne encephalitis (TBE), other lymphocytic meningitis (LMD) and controls (C).

**Table 2.** Specific activity (pKat/µg protein) of lysosomal exoglycosidases (HEX, HEX A, HEX B, GAL, MAN, and GLU) in the urine of patients with tick-borne encephalitis (TBE), other lymphocytic meningitis (LMD), and controls (C).

**Table 3.** HEX A, GAL, MAN and FUC activities expressed per µg creatinine (pKat/µg creatinine) in the urine of patients with tick-borne encephalitis (TBE), other lymphocytic meningitis (LMD) and controls (C).

**Table 4.** Protein and creatinine concentrations in the urine of patients with tick-borne encephalitis (TBE), other lymphocytic meningitis (LMD) and in the control group (C).

## Notes

### Competing Interest Statement

The authors have declared no competing interest.

